# NDST1 as a substrate-reduction target in Mucopolysaccharidosis type IIIC: virtual screening, microsecond molecular dynamics, and peptide design

**DOI:** 10.64898/2026.08.09.743834

**Authors:** Keshav Mohan, Yash Bhargava

**Affiliations:** Bhargava Systems Research Inc, Carmel, IN, USA

**Keywords:** Sanfilippo syndrome, Heparan sulfate, NDST1, Substrate reduction therapy, Molecular docking, Molecular dynamics

## Abstract

Mucopolysaccharidosis IIIC (Sanfilippo syndrome type C) is a rare lysosomal storage disorder caused by loss-of-function mutations in *HGSNAT*, which encodes an enzyme involved in heparan sulfate (HS) degradation, leading to impaired HS catabolism, lysosomal accumulation, and progressive neurodegeneration. Because enzyme replacement therapies have limited penetration across the blood–brain barrier, substrate-reduction therapy represents an alternative therapeutic strategy. Here, N-deacetylase/N-sulfotransferase 1 (NDST1), a key enzyme responsible for HS biosynthesis, was investigated as a potential substrate-reduction target. A structure-based computational pipeline was used to identify and evaluate inhibitors targeting the NDST1 sulfotransferase domain. Approximately 4.1 million drug-like compounds and FDA-approved drugs were screened by molecular docking, followed by pharmacokinetic filtering, molecular dynamics simulations, and MM/PBSA binding free energy calculations. In parallel, peptide binders targeting the same site were generated using diffusion-based protein design and evaluated using molecular dynamics and MM/GBSA analysis. Four chemically distinct small-molecule scaffolds and three peptide candidates were identified as stable binders to the NDST1 active site. The lead small-molecule candidate exhibited a predicted binding free energy of −13.36 ± 5.87 kcal mol^−1^. These provide a focused set of candidates for further investigation and support the feasibility of targeting NDST1 as a substrate-reduction strategy for MPS IIIC.

## 1. Introduction

Sanfilippo syndrome type C (mucopolysaccharidosis IIIC, MPS IIIC) is a rare autosomal recessive lysosomal storage disorder and one of four subtypes of mucopolysaccharidosis type III, each defined by deficiency of an enzyme required for the stepwise lysosomal degradation of heparan sulfate (HS) [1, 2, 3]. MPS IIIC arises from mutations in *HGSNAT*, which encodes heparan–glucosaminide N-acetyltransferase [4, 5]. This lysosomal membrane enzyme transfers an acetyl group to the free amine of terminal glucosamine residues, a modification necessary for the enzymes of the pathway to continue chain degradation [6, 7, 8]. When HGSNAT activity is lost, HS catabolism stalls at this step and undegraded HS accumulates within the lysosome. Most disease-causing missense mutations cause the newly made protein to misfold and become abnormally glycosylated. As a result, the protein is retained and degraded in the endoplasmic reticulum instead of being delivered to the lysosome [9], leading to the cell having no active HGSNAT at the site where the acetylation step must occur.

The consequences of that accumulation are predominantly neurological. HS proteoglycans regulate fibroblast growth factor and Wnt signalling during neurodevelopment, and disturbance of HS structure perturbs the extracellular gradients and receptor interactions through which that regulation operates [10, 11, 12]. Progressive lysosomal engorgement also impairs autophagic flux and weakens postsynaptic signalling. MPS IIIC neurons develop fewer and less mature dendritic spines, along with smaller, less organised postsynaptic densities. Excitatory postsynaptic currents are correspondingly reduced in amplitude, consistent with the weakened excitatory transmission observed in the cortex of MPS IIIA mouse models [13, 14]. In MPS IIIC mouse models, these deficiencies are accompanied by neuroinflammation and progressive neurodegeneration [15, 16]. Infants with MPS IIIC appear unaffected at birth; developmental delay, speech impairment, autism-like behaviour, sleep disturbance and subsequent developmental regression emerge between one and four years of age, followed by motor deterioration and seizures [1, 2].

Current therapies include enzyme replacement, which is effective for the somatic manifestations of several other lysosomal storage disorders. However, it is largely excluded from the central nervous system by the blood–brain barrier and is therefore poorly suited to a primarily neurological indication [17]. This gap motivates substrate reduction, in which biosynthesis of the accumulating macromolecule is partially inhibited so that the residual catabolic capacity of the cell is no longer overwhelmed [18, 19]. Substrate reduction already has direct precedent in the mucopolysaccharidoses, and the existing examples span three distinct ways of slowing the same biosynthetic pathway. The first acts upstream, on the signalling that drives glycosaminoglycan production. Genistein, a plant-derived isoflavone, dampens growth-factor signalling and lowers the amount of stored glycosaminoglycan in skin fibroblasts taken from MPS patients [20]. The second acts more directly on the biosynthetic machinery itself. Rhodamine B, a small dye-like molecule, slows glycosaminoglycan synthesis, and in a mouse model of MPS IIIA this reduces the stored burden and measurably improves behaviour [21, 22]. The third targets the chain-initiating enzymes directly. Silencing EXTL2 and EXTL3, the two enzymes that assemble the HS chain in the first place, likewise reduces HS storage in cells that accumulate it [23]. These examples set a precedent that biosynthesis of an accumulating glycosaminoglycan can be slowed at the level of upstream signalling, of the synthetic machinery itself, or of the chain-initiating enzymes, establishing substrate reduction as a validated strategy for this class of disease rather than a purely theoretical one. Additionally, NDST1 has already been validated as a substrate-reduction therapy target, as transcriptional inhibition of its expression reduces heparan sulfate (HS) sulfation, further supporting the rationale for developing direct NDST1 inhibitors [30].

Prior work has been done with the goal of inhibiting NDST1. A phage-display screen of cyclic ten-residue peptides against the recombinant enzyme returned two inhibitors, CRGWRGEKIGNC and the linear NMQALSMPVT; the first works indirectly, binding HS and blocking the substrate from reaching the enzyme, and only the second contacts the enzyme near its active site [24].

We selected N-deacetylase/N-sulfotransferase 1 (NDST1), the first and rate-influencing modifying enzyme of HS biosynthesis. NDST1 is one of four vertebrate isozymes (NDST1–4) [25] and is a bifunctional type II Golgi membrane protein, in which an N-deacetylase domain removes the acetyl group from N-acetylglucosamine residues of the nascent chain and a sulfotransferase domain then transfers sulfate from 3-phosphoadenosine-5-phosphosulfate (PAPS) to the liberated amine [26, 27]. Because N-sulfation is the initiating modification that licenses all downstream epimerisation and O-sulfation, NDST1 activity sets the overall sulfation density and domain organisation of the mature polysaccharide [10, 28, 29].

The therapeutic rationale follows from the position of the metabolic block. Heavily N-sulfated HS domains place a greater demand on the coordinated set of lysosomal HS-degrading enzymes, and in MPS IIIC every chain must pass through the HGSNAT-catalysed N-acetylation step that the patient can only perform at greatly reduced efficiency. Partial suppression of NDST1 N-sulfotransferase activity lowers the N-sulfation of newly synthesised HS, reduces the degree of sulfation of heparan sulfate chains, and so slows net lysosomal accumulation upstream of the block.

*Ndst1*-null mice die in the perinatal period with pulmonary hypoplasia, craniofacial defects and forebrain hypoplasia [31, 32, 33], whereas heterozygotes carrying a single functional allele are phenotypically normal, indicating that substantial reductions in NDST1 activity are tolerated provided some activity remains [31]. Because complete loss of NDST1 is not tolerated, inhibition will need to be carefully controlled. These constraints underscore the need for selective, structure-guided strategies capable of achieving partial modulation of NDST1 activity.

A cryo-electron microscopy structure of the human NDST1 sulfotransferase domain, together with the rest of the enzyme’s Golgi-facing region (PDB ID 8CCY), has been determined at 2.7 Å resolution [26], and its catalytic donor and acceptor region forms a single large cleft that can be targeted directly, making the domain well suited to structure-based design. In this study, we combined empirical pocket detection, structure-based virtual screening of approximately 4.1 million compounds, ADMET and blood–brain-barrier filtering, microsecond molecular dynamics (MD) simulations and end-point MM/PBSA free-energy calculations to identify and rank candidate NDST1 sulfotransferase-domain ligands, and applied RFdiffusion and ProteinMPNN to generate peptide binders against the same cleft.

## 2. Methods

### 2.1. Protein structure preparation

The structure of human NDST1 (PDB ID 8CCY) was retrieved from the Protein Data Bank [34]. This entry is a single-particle cryo-electron microscopy reconstruction determined at 2.7 Å resolution and covers residues 79– 882 of UniProt P52848, comprising the non-catalytic N-terminal region, the deacetylase domain, and the sulfotransferase domain targeted in this work, together spanning the Golgi-facing part of the enzyme. The cytoplasmic tail and transmembrane anchor (residues 1–78) are not present in the construct, and all simulations reported here therefore describe this Golgi-facing region rather than the intact membrane-embedded enzyme. The deposited structure contains adenosine-3,5-diphosphate (PAP), the desulfated product of the PAPS sulfo-donor, along with bound metal ions and ordered water molecules, all of which were removed prior to docking.

Several residues in the non-catalytic N-terminal and localisation domains were unmodelled, which were built with AlphaFold3 [35], and PDBFixer, distributed as part of the OpenMM package [36]. None of the rebuilt segments forms part of the sulfotransferase catalytic site. All modeled loops lie outside the docking cleft and form no contacts with the ligand, ensuring that the binding site and reported protein–ligand interactions derive entirely from experimentally resolved coordinates. Hydrogen atoms were added at pH 7.4 and the receptor was converted to PDBQT format with Open Babel [37]. Candidate pockets were mapped with the DoGSiteScorer web server [38]. The largest predicted cleft lies within the sulfotransferase domain and coincides with the catalytic donor and acceptor region. The centre of this cleft defined the docking search space.

### 2.2. Ligand libraries and molecular docking

Two libraries were obtained from ZINC [39], approximately 3,000 FDA-approved compounds and approximately

4.1 million natural and drug-like compounds. Structures were desalted, deduplicated, protonated at pH 7.4 and converted from SMILES to PDBQT with Open Babel [37]. Docking was performed with AutoDock Vina [40, 41] using its default scoring function. Ligands were treated as fully flexible and the receptor as rigid. The search space was a 20 × 20 × 20 Å^3^ cube centred on the centroid of the nucleotide bound in the deposited structure (x = 161.24 Å, y = 149.53 Å, z = 127.47 Å), so that the box is anchored on the experimentally observed cofactor position rather than on a predicted site, and exhaustiveness was set to 16. The top-scoring pose of each compound was retained. PAPS was docked under identical conditions as an internal reference; its SMILES string was retrieved from PubChem and prepared by the same protonation and format-conversion route as the library compounds.

### 2.3. ADMET and blood–brain-barrier filtering

Top-ranked compounds were profiled with SwissADME [42]. Because MPS IIIC is a neurological condition, predicted blood–brain-barrier permeability on the BOILED-Egg model was required [43]. Compounds were retained if they satisfied a docking score below −13 kcal mol^−1^ (drug-like library), molecular weight below 500 g mol^−1^, topological polar surface area below 75 Å^2^, fewer than two hydrogen-bond donors, and calculated log P between 1 and 5 (Fig. 2). Retained compounds were additionally screened for PAINS [44, 45] and Brenk [46] structural alerts. For the surviving scaffolds, compliance with Lipinski’s rule of five [47], cytochrome P450 inhibition profile, P-glycoprotein substrate prediction, aqueous solubility and synthetic accessibility were also recorded. Four representative lead classes (Classes 1–4) emerged from the drug-like library, and the blood–brain-barrier-permeable subset of the FDA-approved library was retained for comparison.

**Figure 1:**
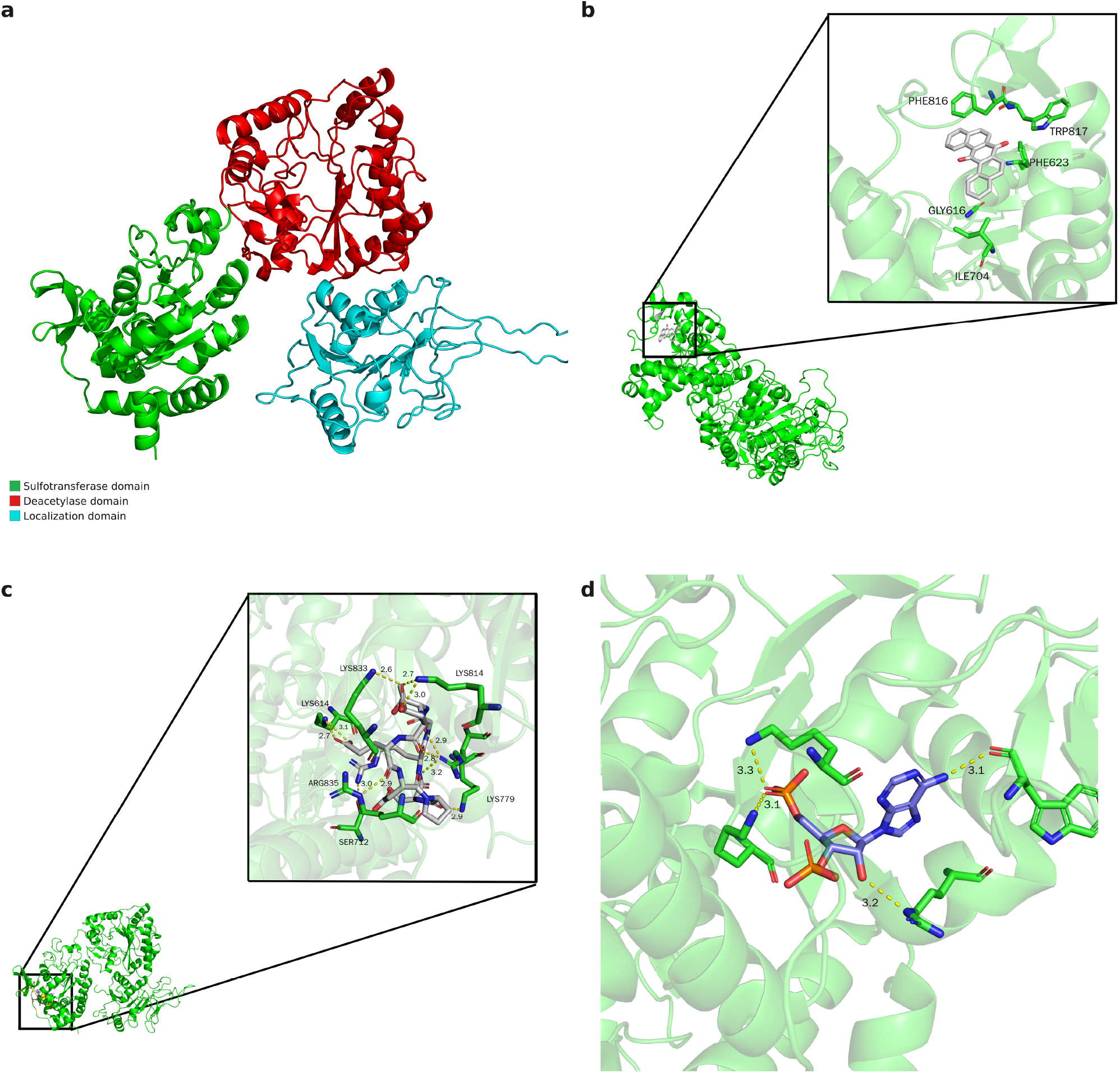
Representative binding poses in the NDST1 sulfotransferase cleft. **a** Domain architecture of the modelled NDST1 construct (residues 79–882, PDB 8CCY): sulfotransferase domain (green), deacetylase domain (red), and localization domain (cyan). **b** The lowest-energy MM/PBSA frame of the lead small-molecule scaffold (Class 2), with contact residues and their distances to the ligand shown (dashed lines; distances in Å). **c** The lowest-energy MM/GBSA frame of Linear Peptide 2 (GTEEDPSR) bound in the cleft, with the basic anchor residues highlighted. **d** The NDST1 sulfotransferase docking pocket (PDB 8CCY) with the deposited nucleotide PAP bound, showing the basic anchor residues that line the cleft and their contact distances to the nucleotide (dashed lines; distances in Å).

**Figure 2:**
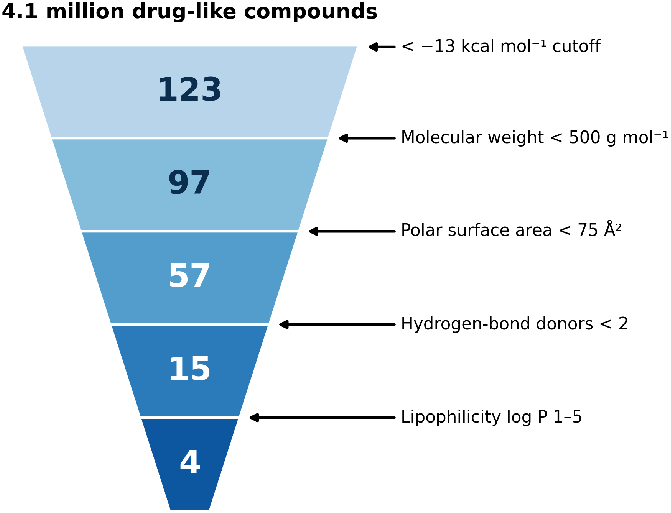
Sequential physicochemical filtering of the 4.1-million-compound small-molecule library, from the initial docking hits down to the four lead scaffolds (Classes 1–4).

### 2.4. Molecular dynamics simulations

Simulations were performed with GROMACS [49] using the CHARMM36 force field for the protein [50], CGenFF parameters for the ligands [51] and the TIP3P water model [52]. Initial ligand poses were generated with Boltz [53]. Each complex was placed in a dodecahedral box with a 1.0 nm solute–boundary buffer, solvated, and neutralised with Na+ and Cl^−^ to a final concentration of 0.15 M. Bonds involving hydrogen were constrained with LINCS [54], permitting a 2 fs integration time step. Long-range electrostatics were treated with the particle-mesh Ewald method [55] using a 1.0 nm real-space cutoff, matched by the van der Waals cutoff. Temperature was maintained at 300 K with velocity rescaling [56] and pressure at 1 bar with the Parrinello–Rahman barostat [57].

Simulations proceeded in two stages. In the initial screening stage, the isolated sulfotransferase domain in complex with each candidate was energy-minimised until the maximum force fell below 500 kJ mol^−1^ nm^−1^, with the backbone restrained, equilibrated for 100 ps in the NVT ensemble and 1 ns in the NPT ensemble, and propagated for 200 ns of unrestrained production dynamics. In the production stage, each ligand was transferred to the complete model, initialised from the final frame of the corresponding screening run, and simulated with the same protocol for 1 *μ*s. The protein (PDB ID 8CCY) with resolved loops and PAP removed was simulated as a control. Coordinates were written every 10 ps.

All trajectories were corrected for periodic boundary conditions before analysis. Ligand root-mean-square deviation (RMSD) was then computed after least-squares fitting each frame to the protein. Per-residue C*α* RMSF was computed over all 804 residues of the receptor. Radius of gyration (Rg) and solvent-accessible surface area (SASA) were computed on the corrected trajectories.

### 2.5. Binding free-energy calculations

End-point binding free energies were computed with gmx_MMPBSA [58], which implements the MMPBSA.py methodology [59] for GROMACS trajectories, using a single-trajectory MM/PBSA scheme with implicit solvent. The binding free energy was decomposed as

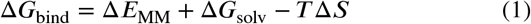

where Δ*E*_MM_ is the gas-phase molecular-mechanics interaction energy (van der Waals and electrostatic terms), Δ*G*_solv_ is the sum of polar and non-polar solvation contributions, and −*T* Δ*S* is the solute entropic penalty. For the four small-molecule classes the polar term was obtained from the Poisson–Boltzmann equation, and both the enthalpic and the entropic terms were evaluated, the latter over the final 250 ns of each 1 *μ*s trajectory.

### 2.6. Peptide binder design

Peptide binders targeting the sulfotransferase-domain cleft of NDST1 were designed using an RFdiffusion– ProteinMPNN pipeline. RFdiffusion [60] generated peptide-binding backbone conformations complementary to the target pocket, and ProteinMPNN [61] assigned amino acid sequences to each designed backbone. A total of 960 candidate sequences were generated and filtered using a ProteinMPNN mean negative log-likelihood (NLL) threshold of <0.8, yielding 48 high-confidence designs. These sequences were clustered into seven sequence families, and one representative from each cluster was evaluated using AlphaFold. Only three clusters produced representatives with an interface predicted TM-score (ipTM) greater than 0.6. The remaining sequences within these three clusters were subsequently predicted with AlphaFold, and the highest-confidence design from each cluster was selected for molecular dynamics simulations.

The three selected binders comprised two linear peptides (Linear Peptide 1, KVWEGIEKRE; Linear Peptide 2, GTEEDPSR, shown in Fig. S1b) and one cyclic peptide (Cyclic Peptide 1, LVERRKED). Each peptide– NDST1 complex was initially simulated for 250 ns using the sulfotransferase-domain molecular dynamics protocol described above; Linear Peptide 2, which remained engaged with the pocket, was extended to a full 1 *μ*s production run, and all Linear Peptide 2 results reported below use this extended trajectory, while Linear Peptide 1 and Cyclic Peptide 1 dissociated within the initial 250 ns and were not extended further. Complex stability was assessed using per-residue C*α* RMSF calculated separately for the receptor and peptide, as peptides that dissociated from the binding pocket exhibited uniformly elevated fluctuations across all residues, whereas stably bound peptides displayed minimal RMSF values, representing localized flexibility. Binding enthalpies were subsequently estimated using the MM/GBSA approach described above, including generalized Born electrostatic and solvent-accessible surface area nonpolar terms. For the peptide complexes, binding enthalpies were computed for all three designs with MM/GBSA, in which the polar solvation term is the generalised-Born energy and the non-polar term is a surface-area (ESURF) contribution. Linear Peptide 1 and Cyclic Peptide 1 were analysed over their full 250 ns trajectories (250 frames) and Linear Peptide 2 over its full 1 *μ*s trajectory (1000 frames), all at 298.15 K with receptor mask :1–804 and ligand mask :805–812, and only the enthalpic contribution was computed in every case; no entropy term was evaluated.

## 3. Results and discussion

### 3.1. The NDST1 sulfotransferase domain presents a large, accessible cleft

DoGSiteScorer identified the largest cleft of the sulfotransferase domain as the primary candidate site (Fig. 1d). The pocket is well within the size range that accommodates drug-like ligands, and coincides with the donor and acceptor region of the catalytic site, with the nucleotide occupying the largest subpocket in the deposited structure, so that a ligand retained here is positioned to occlude catalysis directly.

### 3.2. Docking of the FDA-approved library

To identify potential drug repurposing candidates, we docked an approximately 3,000-compound library of FDA-approved drugs, yielding several high-scoring candidates. Ergotamine (ZINC000052955754) was the highest-ranked at −13.01 kcal mol^−1^, and the mean score across the top 20 compounds was −11.29 kcal mol^−1^. Ergotamine was nevertheless excluded during the filtering stage because it is a nonspecific binder, and its high molecular weight and large polar surface area are incompatible with efficient passive penetration of the central nervous system. Applying the permeability and drug-likeness filters to the top 20 left four library entries in final consideration, each of which scored more favourably than the PAPS reference (−10.23 kcal mol^−1^). These were progesterone (ZINC000004428529, −11.47 kcal mol^−1^), palonosetron, which is present in the library as two separate entries and so accounts for two of the four (ZINC000006094354, listed under the trade name Aloxi, −11.25 kcal mol^−1^; ZINC000003795819, −10.99 kcal mol^−1^), and nebivolol (ZINC000001999441, −10.91 kcal mol^−1^) — three distinct drugs, chemically unrelated to HS biosynthesis; these scores are best read as a ranking of pocket complementarity rather than as evidence of activity. Only 4 of the top 20 entries survived blood–brain-barrier filtering.

### 3.3. Virtual screening of the drug-like library

Docking of approximately 4.1 million drug-like ZINC compounds returned 123 molecules scoring below −13 kcal mol^−1^. Sequential physicochemical filtering then reduced this set stepwise (Fig. 2): a molecular-weight cutoff below 500 g mol^−1^ retained 97 compounds (mean docking score −13.23 kcal mol^−1^), a polar-surface-area cutoff below 75 Å^2^ retained 57, a hydrogen-bond-donor cutoff below 2 retained 15, and a lipophilicity window of log P 1–5 retained 4. These four survivors define the representative lead scaffolds designated Classes 1–4 (Table 1; chemical structures in Fig. S1a), each scoring more favourably than the PAPS reference.

**Table 1.** Representative lead scaffolds identified from the drug-like library, with SMILES and docking scores.

| Class | SMILES | Docking score (kcal mol<sup>-1</sup>) |
| --- | --- | --- |
| 1 | O=C1c2ccccc2C(=O)c2cc3c(cc21)[C@H]1CC[C@H]3C1 | -13.42 |
| 2 | O=C1c2ccc3ccccc3c2C(=O)c2c1ccc1ccccc21 | -13.20 |
| 3 | c1ccc2c(C[C@@H]3COC[C@@H]3NC3CCN(c4ccc5c(c4)CCC5)CC3)ccnc2c1 | -13.02 |
| 4 | Cc1cccc(N2C(=O)NC(=O)/C(=C3cccn3-c3ccc4ccccc4c3)C2=O)c1 | -13.26 |

### 3.4. Predicted ADMET properties and structural liabilities

Predicted physicochemical and pharmacokinetic properties for the four scaffolds are given in Table S1. All four satisfy Lipinski’s rule of five [47] and all return a bioavailability score of 0.55. Class 1 is small and conformationally rigid (MW 274 g mol^−1^, TPSA 34.1 Å^2^, no rotatable bonds, log P 3.16), a profile consistent with hydrophobic contacts and with favourable passive absorption. Class 2 is larger and fully aromatic (MW 308 g mol^−1^, log P 4.41), and its higher lipophilicity is expected to limit aqueous solubility. Class 3 is the most flexible of the set (MW 428 g mol^−1^, five rotatable bonds, log P 4.45) and carries a hydrogen-bond donor, allowing directional polar contacts. Class 4 has highest polar surface area of the set (MW 421 g mol^−1^, TPSA 71.4 Å^2^, log P 3.66), and its size and polarity may reduce membrane and BBB permeability.

Predicted gastrointestinal absorption varied across the set, high for Classes 1 and 2, moderate for Class 3, and reduced for Class 4. All four were nonetheless predicted to be blood–brain-barrier permeant on the BOILED-Egg model [43]. P-glycoprotein prediction flagged Classes 1 and 3 as possible efflux substrates and Classes 2 and 4 as non-substrates; measured permeability, efflux handling and brain exposure remain to be determined. Cytochrome P450 profiling indicated isoform-specific inhibition (Class 1 CYP1A2 and CYP2D6; Class 2 CYP1A2 and CYP2C19; Class 3 CYP2D6 and CYP3A4; Class 4 CYP2C19 and CYP2C9), implying compound-dependent and, manageable drug-interaction risk. Predicted aqueous solubility ranged from moderate to poor across the ESOL, Ali and SILICOS-IT models, as expected for hydrophobic scaffolds, and synthetic accessibility scores of 2.49–4.02 indicate moderate synthetic feasibility.

Structural-alert screening identified a few potential liabilities. Classes 1 and 2 are fused polycyclic quinones, aromatic ring systems carrying a pair of ring ketones, an arrangement that accepts and releases electrons readily and is therefore chemically reactive within the body. Quinones are among the most frequently flagged pan-assay-interference chemotypes and are a recognised source of false-positive readouts through redox cycling, covalent reactivity, metal chelation and colloidal aggregation [44, 45]. Classes 3 and 4 remain the cleaner starting points, though Class 4 also carries isolated Brenk alerts [46], and we prioritise them accordingly.

### 3.5. Predicted binding modes

Interaction profiling of the docked poses with PLIP [48] revealed a common anchoring pattern across the four scaffolds, together with a clear gradient in the number of contacts each scaffold makes (Fig. 3).

**Figure 3:**
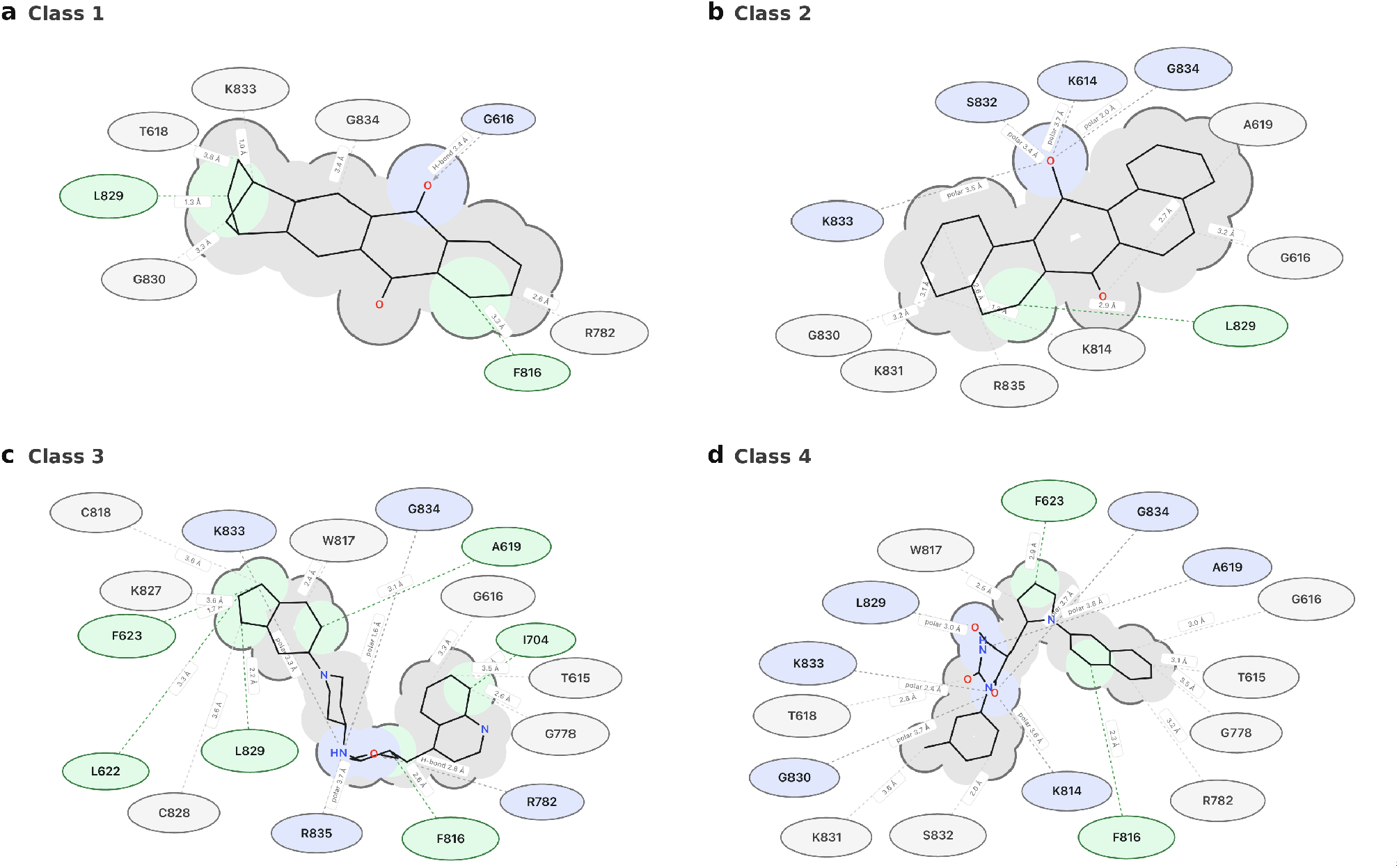
PLIP-predicted binding modes of the four small-molecule lead scaffolds in the NDST1 pocket (**a** Class 1, **b** Class 2, **c** Class 3, **d** Class 4).

Class 1, the smallest and most rigid scaffold, engages the fewest residues (Fig. 3a), forming a single hydrogen bond from one quinone carbonyl to Gly616, hydrophobic contacts to Leu829 and Phe816, and further contacts to Thr618, Lys833, Gly830, Gly834 and Arg782. Class 2 presents both carbonyls to the polar face of the pocket (Fig. 3b), forming polar contacts with Lys614, Ser832, Gly834 and Lys833, a hydrophobic contact to Leu829, and additional contacts to Ala619, Gly616, Gly830, Lys831, Arg835 and Lys814. Class 3 forms by far the most extensive network of the set (Fig. 3c), consistent with its greater size and flexibility, forming a hydrogen bond to the guanidinium of Arg782, polar contacts to Gly834, Arg835 and Lys833, an aromatic and hydrophobic shell comprising Phe623, Phe816, Ala619, Leu622, Leu829 and Ile704, and further contacts to Cys818, Trp817, Lys827, Cys828, Thr615, Gly778 and Gly616. Class 4 spans the cleft with its polar core facing the basic rim (Fig. 3d), making polar contacts with Lys833, Leu829, Gly830, Gly834, Ala619 and Lys814, aromatic contacts with Phe623 and Phe816, and further contacts to Trp817, Thr618, Lys831, Ser832, Gly616, Thr615, Gly778 and Arg782.

Arg782, Gly834, Lys833, Phe816, Leu829 and Ala619 recur across all four classes and constitute the principal anchor set of this pocket, with Gly616, Gly830 and Ser832 appearing in three of the four. This set overlaps with the cofactor site annotated for NDST1 in UniProt P52848, which lists residues 614–618, 712, 817 and 833–837 as nucleotide-binding and identifies Lys614 as the catalytic residue for sulfotransferase activity. The scaffolds therefore occupy the donor subpocket and engage the catalytic lysine itself, which is the structural basis for the expectation that a ligand retained here would occlude sulfuryl transfer. The aromatic pair Phe623 and Phe816 provides a nonpolar contact surface that Class 3 and Class 4 exploit most fully, while Classes 1 and 2 rely more on the polar rim.

### 3.6. The scaffolds remain bound over microsecond simulations

Across 1 *μ*s of production dynamics all four scaffolds remained bound in the sulfotransferase site, and none dissociated (Fig. 4). Ligand RMSD, which reports displacement within the site, plateaued at 0.451 ± 0.019 Å for Class 1 and 0.474 ± 0.014 Å for Class 2, and at a higher but equally stable plateau of 0.769 ± 0.012 Å for Class 3 and 0.708 ± 0.010 Å for Class 4 (mean ± block standard error over 20 ns blocks; Fig. 4a). Every system reached its plateau within approximately the first 100 ns and held it thereafter, and in no run did the ligand exceed 1.2 Å of displacement from its fitted pose.

**Figure 4:**
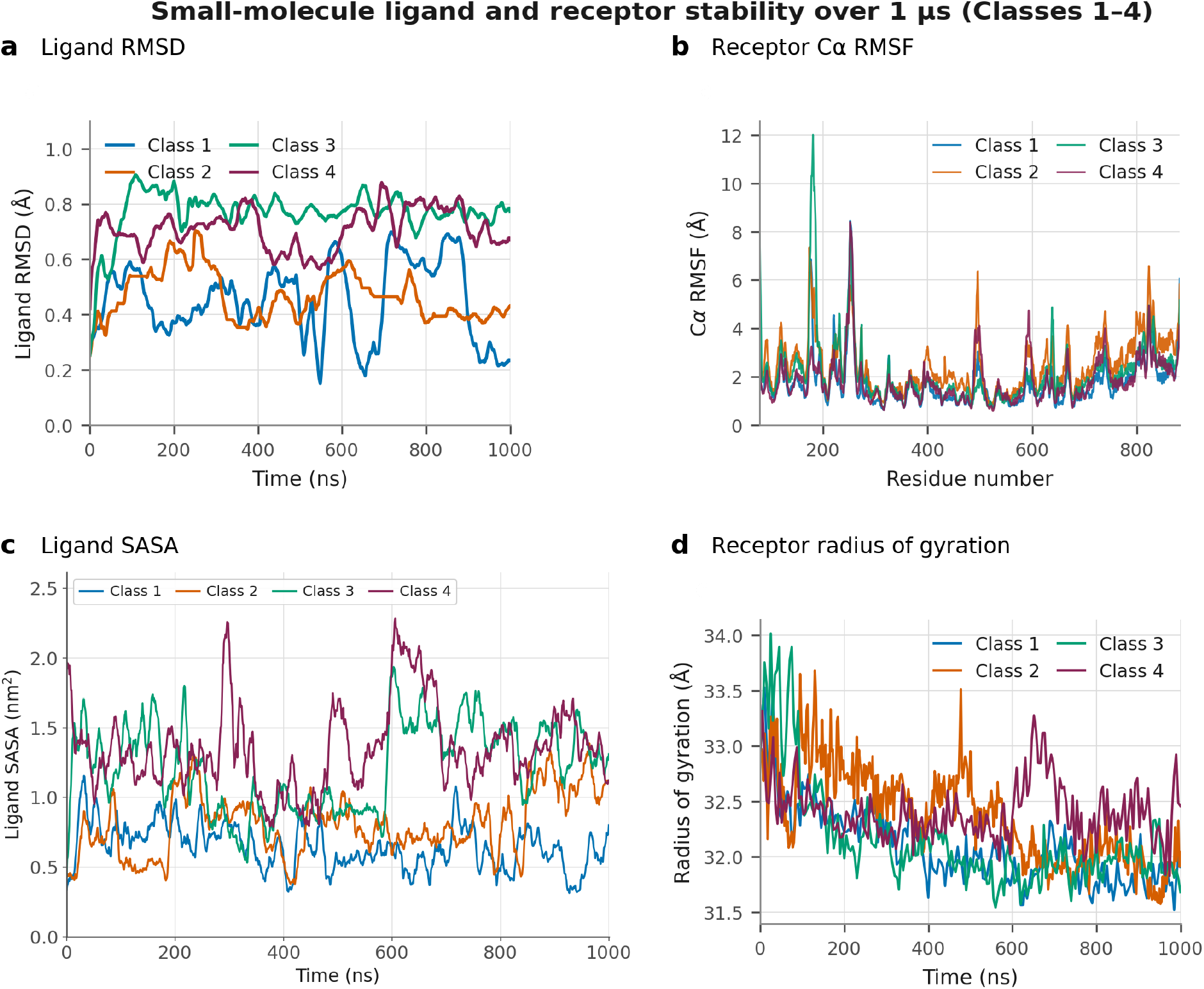
Small-molecule complex stability over the 1 µs production trajectories (Classes 1–4). **a** Ligand RMSD. **b** Receptor C*α* RMSF. **c** Ligand SASA. **d** Receptor radius of gyration.

Per-residue C*α* RMSF profiles were closely superimposable across the four complexes throughout the structured core, remaining within approximately 1–2 Å and rising only at solvent-exposed loops and at the termini, as expected for a folded catalytic domain (Fig. 4b). Whole-protein mean C*α* RMSF values were 1.74, 2.49, 2.15 and 1.84 Å for Classes 1–4, with Class 1 the most rigid. Radius of gyration remained tightly confined in every complex, with per-run means of 32.09 ± 0.35, 32.37 ± 0.43, 32.15 ± 0.47 and 32.41 ± 0.26 Å for Classes 1–4 and a standard deviation below 0.5 Å throughout (Fig. 4d). All four systems contracted by between 0.5 and 1.1 Å from their starting geometry as the modelled loops relaxed, after which Rg fluctuated about a stationary value and the four complexes became indistinguishable within those fluctuations, indicating that ligand binding does not perturb the compact fold.

Ligand-exposed surface area remained small in every case and increased modestly across the series (0.65 ± 0.24, 0.81 ± 0.30, 1.22 ± 0.39 and 1.36 ± 0.42 nm^2^ for Classes 1–4; Fig. 4c), confirming that each ligand stayed enclosed within the pocket. The ordering of ligand SASA tracks the ordering of ligand RMSD, with the most rigidly held scaffolds presenting the least solvent exposure. All four ligands remained associated with the pocket throughout the simulations.

### 3.7. MM/PBSA ranking identifies Class 2 as the strongest predicted binder

End-point MM/PBSA calculations resolved binding-strength differences that docking had not (Table 2, Fig. 5). All four ligands were enthalpically favourable, with combined gas-phase and solvation enthalpies of −15.87 ± 0.20, −22.68 ± 0.34, −19.37 ± 0.30 and −22.09 ± 0.36 kcal mol^−1^ for Classes 1–4, and entropic penalties (−*T* Δ*S*) ranging from 7.42 to 13.06 kcal mol^−1^. Combining the terms placed Class 2 strongest at −13.36 ± 5.87 kcal mol^−1^, with Class 3 (−11.96 ± 3.06 kcal mol^−1^) and Class 4 (−11.90 ± 3.52 kcal mol^−1^) essentially indistinguishable just behind it, and Class 1 markedly weaker at −2.81 ± 2.77 kcal mol^−1^. The overall ranking is Class 2 > Class 3 ≈ Class 4 > Class 1. In every complex binding is driven by the van der Waals term and opposed by polar desolvation, with the electrostatic and non-polar solvation contributions smaller in magnitude (Table 2). This is the signature of a predominantly shape-complementary, hydrophobically driven interaction in a solvent-exposed groove.

**Table 2.** MM/PBSA binding free energies and energy components for the four lead scaffolds (kcal mol^−1^)

| Class | $\Delta E_{\text{vdW}}$ | $\Delta E_{\text{elec}}$ | $\Delta G_{\text{polar}}$ | Enthalpy ($\Delta H$) | $-T\Delta S$ | $\Delta G_{\text{bind}}$ |
| --- | --- | --- | --- | --- | --- | --- |
| 1 | -31.04 | -13.38 | 31.84 | $-15.87 \pm 0.20$ | $13.06 \pm 0.01$ | $-2.81 \pm 2.77$ |
| 2 | -37.29 | -14.12 | 32.38 | $-22.68 \pm 0.34$ | $9.32 \pm 0.94$ | $-13.36 \pm 5.87$ |
| 3 | -41.99 | -4.03 | 31.64 | $-19.37 \pm 0.30$ | $7.42 \pm 0.01$ | $-11.96 \pm 3.06$ |
| 4 | -44.35 | -8.35 | 35.37 | $-22.09 \pm 0.36$ | $10.19 \pm 0.01$ | $-11.90 \pm 3.52$ |

**Figure 5:**
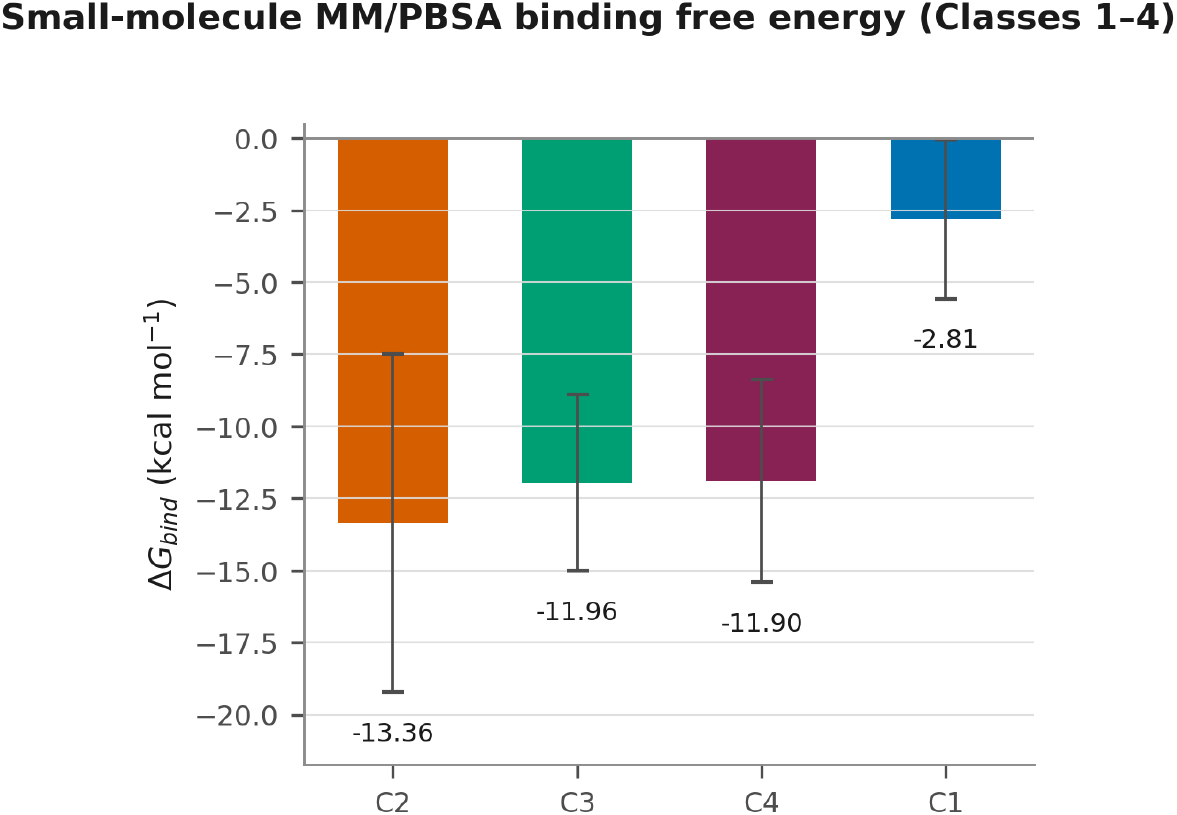
End-point MM/PBSA ranked total binding free energy of the four small-molecule scaffolds (Classes 1–4, with SEM; see Table 2 for energy components).

The binding free-energy analysis reveals several features that are not apparent from docking results alone. Class 1 achieved the most favourable docking score of the four (−13.42 kcal mol^−1^), yet exhibited by far the least favourable MM/PBSA binding free energy. In contrast, Class 2 ranked only third by docking score (−13.20 kcal mol^−1^) and displayed a comparatively modest PLIP contact network (above), but yielded the most favourable end-point free energy. Per-residue energy decomposition (Fig. 7b; component breakdown in Fig. S2b) attributes this behaviour to an unusually strong anchoring interaction, with Phe816 contributing −1.79 kcal mol^−1^ to Class 2—the largest single stabilising per-residue contribution observed among all four complexes and arising almost entirely from van der Waals packing. Moreover, the difference between Class 3 and Class 4 (−11.96 ± 3.06 versus −11.90 ± 3.52 kcal mol^−1^) is substantially smaller than the uncertainty associated with either estimate, supporting only the conclusion that Classes 2, 3, and 4 all bind more favourably than Class 1 without establishing a definitive rank order among the top three candidates. The entropic penalties for Classes 1, 3, and 4 also exhibit markedly smaller dispersions (0.01 kcal mol^−1^ each) than that of Class 2 (0.94 kcal mol^−1^).

### 3.8. Per-residue MM-PBSA decomposition of the four lead scaffolds

Decomposing each complex’s MM-PBSA free energy into per-residue contributions resolves the residue-level origin of the ranking established above (Table 2, Fig. 7; component breakdown in Fig. S2). In every complex a small set of hydrophobic and aromatic residues dominates the favourable binding energy, driven overwhelmingly by van der Waals packing, with electrostatics contributing comparatively little and, for several residues, polar desolvation actively opposing the interaction — the same pattern seen at the level of the whole complex (Table 2). Phe816 is the single most consistent stabiliser, contributing Δ*G*_total_ of −0.98, −1.79, −0.69 and −1.50 kcal mol^−1^ in Classes 1–4 respectively (Fig. 7a–d), and a recurring cluster of residues (Ala619, Gly616, Leu622, Phe623, Ile704 and Leu781) lines the pocket across all four complexes, in agreement with the anchor residues identified independently by PLIP contact profiling above. Class 4’s tightest single stabiliser is Leu829 (−2.16 kcal mol^−1^) and Class 3 additionally recruits Ile704 and Tyr837, whereas Class 1’s profile is the most diffuse of the four, with no single residue contributing below −1.0 kcal mol^−1^ (Fig. 7a), consistent with its markedly lower overall free energy (Fig. 5).

### 3.9. Peptide binders

Three designs generated by the RFdiffusion– ProteinMPNN pipeline were carried into 250 ns simulations; Linear Peptide 2 was subsequently extended to a full 1 *μ*s production run (see below) (Table S2, Fig. 6). In all three complexes the NDST1 receptor remained folded and compact (mean receptor C*α* RMSF 2.56 Å for Linear Peptide 1, 1.78 Å for Cyclic Peptide 1 and 2.24 Å for Linear Peptide 2; radius of gyration 32.94 ± 0.43, 32.67 ± 0.27 and 32.92 ± 0.39 Å respectively, standard deviation below 0.5 Å throughout), so peptide binding left the receptor fold intact (Fig. 6b).

**Figure 6:**
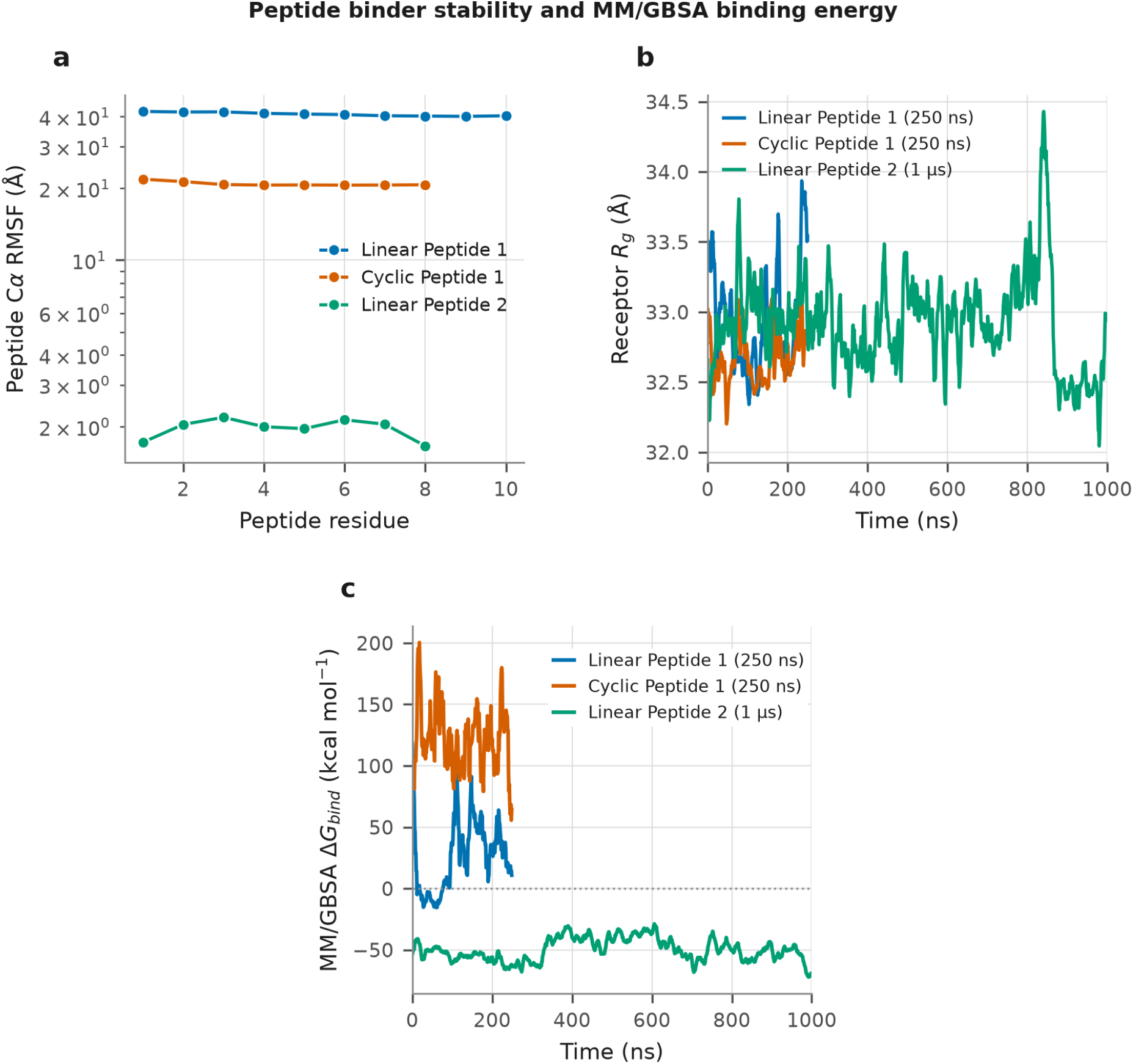
Stability and binding energetics of the three RFdiffusion/ProteinMPNN peptide binders (Linear Peptide 2 over its full 1 *μ*s trajectory; Linear Peptide 1 and Cyclic Peptide 1 over 250 ns). **a** Peptide C*α* RMSF. **b** Receptor radius of gyration. **c** MM/GBSA binding free energy over time (see Table 3 for energy components).

**Figure 7:**
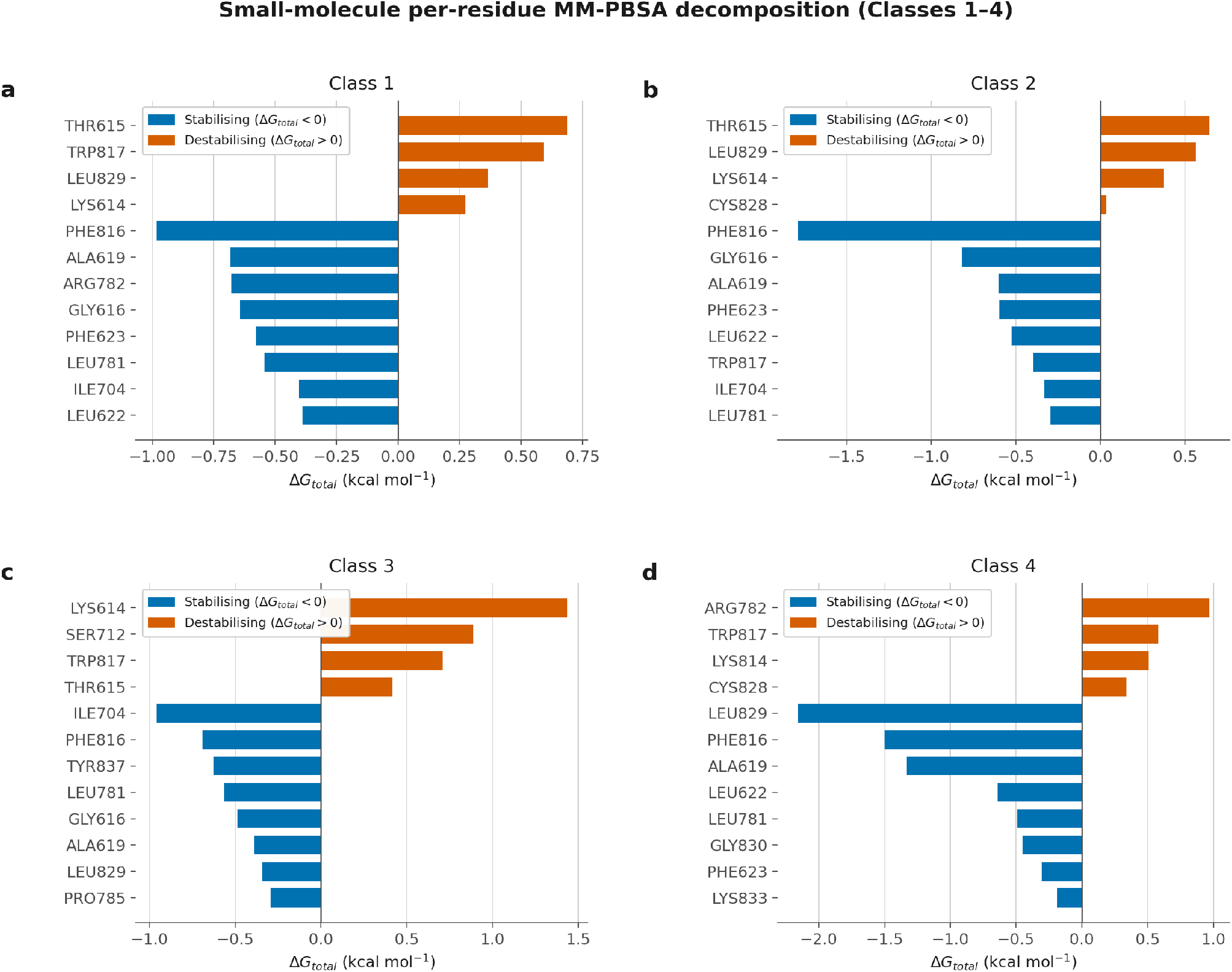
Per-residue MM-PBSA decomposition for the four small-molecule scaffolds, showing the residues that most stabilise (blue) or destabilise (orange) binding in each class (**a**–**d**, Classes 1–4).

Retention within the pocket differed sharply between designs. Only Linear Peptide 2 (GTEEDPSR) remained engaged, and was therefore extended from the initial 250 ns screen to a full 1 *μ*s production run: across that trajectory its peptide C*α* RMSF stayed low (1.66–2.19 Å per residue, mean 1.97 Å), comparable to the internal flexibility of a folded surface loop, the receptor fold remained intact (Rg 32.92 ± 0.39 Å), and the distance from the peptide centre of mass to the basic anchor residues stayed stable at 14.1 ± 0.56 Å (range 12.8–16.0 Å), indicating sustained engagement with the pocket rather than transient rebinding. Cyclic Peptide 1 and Linear Peptide 1, each simulated for 250 ns, both showed large, near-uniform per-residue displacements (20.9 ± 0.42 Å and 40.94 ± 0.71 Å; Fig. 6a) characteristic of whole-peptide rigid-body motion, indicating that neither remained bound at the site and so were not extended further.

**Table 3.**
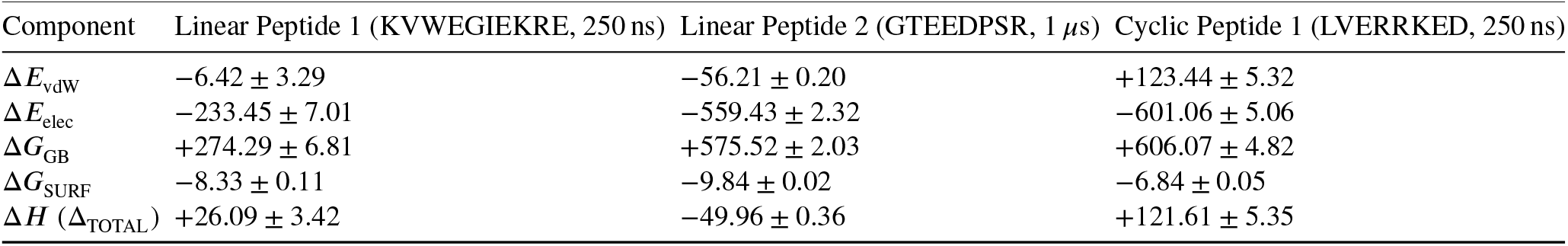
MM/GBSA energy decomposition for the three peptide complexes (kcal mol^−1^, enthalpic terms only) Mean ± SEM over the full simulated trajectory for each design.

Binding enthalpies were computed for all three designs by MM/GBSA (Table 3; Fig. 6c). The van der Waals term is the primary driver of the difference. Over its full 1 *μ*s trajectory, Linear Peptide 2 has a large attractive ΔE_vdW of −56.21 ± 0.20 kcal mol^−1^, indicating extensive and persistent surface contact, whereas the other two designs, simulated for 250 ns, show a near-vanishing or strongly positive van der Waals term (Linear Peptide 1, −6.42 ±3.29; Cyclic Peptide 1, +123.44 ± 5.32 kcal mol^−1^), consistent with dissociation from the site.

Only Linear Peptide 2 gives a favourable total, at −49.96 ± 0.36 kcal mol^−1^ by MM/GBSA over its full 1 *μ*s trajectory. Its interface is driven by strong electrostatics (−559.43 ± 2.32 kcal mol^−1^) and van der Waals contact, largely but not entirely offset by the polar desolvation penalty (+575.52 ± 2.03 kcal mol^−1^). For Linear Peptide 2, the binding free energy remains negative and stable throughout the trajectory, whereas Linear Peptide 1 and Cyclic Peptide 1 fluctuate around substantially positive values.

Only Linear Peptide 2 therefore remained bound in the active site (Fig. 1c). Its sequence complements the basic anchor set (Arg782, Lys614, Lys833, Lys814) identified in the small-molecule binding modes and points to electrostatic complementarity as the dominant determinant of recognition at this site.

## 4. Conclusion

This study maps an open, druggable catalytic cleft on the NDST1 sulfotransferase domain, screens approximately 4.1 million compounds against it, and carries four filtered small-molecule scaffolds and three peptide designs through molecular dynamics and end-point free-energy analysis. All four scaffolds remained bound in the sulfotransferase site over 1 *μ*s without perturbing the receptor fold, and MM/PBSA ranked them Class 2 > Class 3 ≈ Class 4 > Class 1, with Class 2 the strongest predicted binder at −13.36 ± 5.87 kcal mol^−1^ (Classes 3 and 4 statistically indistinguishable from it and from each other). Because Classes 1 and 2 are quinones with recognised pan-assay-interference liabilities, Classes 3 and 4 represent the cleaner starting points despite Class 2’s favourable computed energetics. Of the peptide designs, only the linear peptide GTEEDPSR remained engaged, returning an MM/GBSA binding enthalpy of −49.96 ± 0.36 kcal mol^−1^ over its full 1 *μ*s trajectory, where the other two designs, simulated for 250 ns, gave large positive values; its sequence complements the basic anchor set of Arg782, Lys614, Lys833 and Lys814 that recurs across the small-molecule binding modes, identifying electrostatic complementarity as the principal determinant of recognition at this site. Next steps involve in vitro measurement of NDST1 sulfotransferase inhibition for Classes 3 and 4, together with the interference controls required to clear the quinone scaffolds. Additionally, counter-screening against NDST2–4 to establish isozyme selectivity, since a compound that cannot discriminate among the isozymes is disqualified regardless of potency, and then cellular assays of HS sulfation and lysosomal storage in patient-derived MPS IIIC fibroblasts and neurons.

## Funding

The authors declare that no funds, grants, or other support were received during the preparation of this manuscript.

## Declaration of competing interest

The authors have no relevant financial or non-financial interests to disclose.

## CRediT authorship contribution statement

**Keshav Mohan**: Conceptualization, Methodology, Software, Formal analysis, Investigation, Data curation, Writing – original draft, Writing – review & editing. **Yash Bhargava**: Conceptualization, Methodology, Software, Formal analysis, Investigation, Data curation, Writing – original draft, Writing – review & editing.

## Ethical approval

Not applicable. This study involved no human participants, animal subjects or patient-derived material.

## Data availability

Molecular-dynamics trajectories, analysis scripts and the compiled time-series data supporting the findings of this study are available from the corresponding author upon request.

## Notes

### Competing Interest Statement

The authors have declared no competing interest.

## References

[1] Heon-Roberts R, Nguyen ALA, Pshezhetsky AV (2020) Molecular bases of neurodegeneration and cognitive decline, the major burden of Sanfilippo disease. J Clin Med 9:344.

[2] Valstar MJ, Ruijter GJG, van Diggelen OP, Poorthuis BJ, Wijburg FA (2008) Sanfilippo syndrome: a mini-review. J Inherit Metab Dis 31:240–252.

[3] Muschol N, Giugliani R, Jones SA, Muenzer J, Smith NJC, Whitley CB, Donnell M, Drake E, Elvidge K, Melton L, O’Neill C (2022) Sanfilippo syndrome: consensus guidelines for clinical care. Orphanet J Rare Dis 17:391.

[4] Hrebicek M, Mrazova L, Seyrantepe V, Durand S, Roslin NM, Noskova L, Hartmannova H, Ivanek R, Cizkova A, Poupetova H, Sikora J, Urinovska J, Stranecky V, Zeman J, Lepage P, Roquis D, Verner A, Ausseil J, Beesley CE, Maire I, Poorthuis BJ, van de Kamp J, van Diggelen OP, Wevers RA, Hudson TJ, Fujiwara TM, Majewski J, Morgan K, Kmoch S, Pshezhetsky AV (2006) Mutations in TMEM76* cause mucopolysaccharidosis IIIC (Sanfilippo C syndrome). Am J Hum Genet 79:807–819.

[5] Fan X, Zhang H, Zhang S, Bagshaw RD, Tropak MB, Callahan JW, Mahuran DJ (2006) Identification of the gene encoding the enzyme deficient in mucopolysaccharidosis IIIC (Sanfilippo disease type C). Am J Hum Genet 79:738–744.

[6] Durand S, Feldhammer M, Bonneil E, Thibault P, Pshezhetsky AV (2010) Analysis of the biogenesis of heparan sulfate acetyl-CoA:-glucosaminide N-acetyltransferase provides insights into the mechanism underlying its complete deficiency in mucopolysaccharidosis IIIC. J Biol Chem 285:31233–31242.

[7] Fan X, Tkachyova I, Sinha A, Rigat B, Mahuran D (2011) Characterization of the biosynthesis, processing and kinetic mechanism of action of the enzyme deficient in mucopolysaccharidosis IIIC. PLoS One 6:e24951.

[8] Navratna V, Kumar A, Rana JK, Mosalaganti S (2024) Structure of the human heparan–glucosaminide N-acetyltransferase (HGSNAT). eLife 13:RP93510.

[9] Feldhammer M, Durand S, Pshezhetsky AV (2009) Protein misfolding as an underlying molecular defect in mucopolysaccharidosis III type C. PLoS One 4:e7434.

[10] Esko JD, Selleck SB (2002) Order out of chaos: assembly of ligand binding sites in heparan sulfate. Annu Rev Biochem 71:435–471.

[11] Kjellén L, Lindahl U (2018) Specificity of glycosaminoglycan– protein interactions. Curr Opin Struct Biol 50:101–108.

[12] Pan Y, Woodbury A, Esko JD, Grobe K, Zhang X (2006) Heparan sulfate biosynthetic gene Ndst1 is required for FGF signaling in early lens development. Development 133:4933–4944.

[13] Para C, Bose P, Bruno L, Freemantle E, Taherzadeh M, Pan X, Han C, McPherson PS, Lacaille JC, Bonneil E, Thibault P, O’Leary C, Bigger B, Morales CR, Di Cristo G, Pshezhetsky AV (2021) Early defects in mucopolysaccharidosis type IIIC disrupt excitatory synaptic transmission. JCI Insight 6:e142073.

[14] Dwyer CA, Scudder SL, Lin Y, Dozier LE, Phan D, Allen NJ, Patrick GN, Esko JD (2017) Neurodevelopmental changes in excitatory synaptic structure and function in the cerebral cortex of Sanfilippo syndrome IIIA mice. Sci Rep 7:46576.

[15] Settembre C, Fraldi A, Jahreiss L, Spampanato C, Venturi C, Medina D, de Pablo R, Tacchetti C, Rubinsztein DC, Ballabio A (2008) A block of autophagy in lysosomal storage disorders. Hum Mol Genet 17:119–129.

[16] Martins C, Hulkova H, Dridi L, Dormoy-Raclet V, Grigoryeva L, Choi Y, Langford-Smith A, Wilkinson FL, Ohmi K, DiCristo G, Hamel E, Ausseil J, Cheillan D, Moreau A, Svobodova E, Hajkova Z, Tesarova M, Hansikova H, Bigger BW, Hrebicek M, Pshezhetsky AV (2015) Neuroinflammation, mitochondrial defects and neurodegeneration in mucopolysaccharidosis III type C mouse model. Brain 138:336–355.

[17] Pardridge WM (2012) Drug transport across the blood–brain barrier. J Cereb Blood Flow Metab 32:1959–1972.

[18] Coutinho MF, Santos JI, Alves S (2016) Less is more: substrate reduction therapy for lysosomal storage disorders. Int J Mol Sci 17:1065.

[19] Jakóbkiewicz-Banecka J, Węgrzyn A, Węgrzyn G (2007) Substrate deprivation therapy: a new hope for patients suffering from neuronopathic forms of inherited lysosomal storage diseases. J Appl Genet 48:383–388.

[20] Piotrowska E, Jakóbkiewicz-Banecka J, Barańska S, Tylki-Szymańska A, Czartoryska B, Węgrzyn A, Węgrzyn G (2006) Genistein-mediated inhibition of glycosaminoglycan synthesis as a basis for gene expression-targeted isoflavone therapy for mucopolysaccharidoses. Eur J Hum Genet 14:846–852.

[21] Roberts AL, Thomas BJ, Wilkinson AS, Fletcher JM, Byers S (2006) Inhibition of glycosaminoglycan synthesis using rhodamine B in a mouse model of mucopolysaccharidosis type IIIA. Pediatr Res 60:309–314.

[22] Roberts AL, Rees MH, Klebe S, Fletcher JM, Byers S (2007) Improvement in behaviour after substrate deprivation therapy with rhodamine B in a mouse model of MPS IIIA. Mol Genet Metab 92:115–121.

[23] Kaidonis X, Liaw WC, Roberts AD, Ly M, Anson D, Byers S (2010) Gene silencing of EXTL2 and EXTL3 as a substrate deprivation therapy for heparan sulphate storing mucopolysaccharidoses. Eur J Hum Genet 18:194–199.

[24] Gesteira TF, Coulson-Thomas VJ, Taunay-Rodrigues A, Oliveira V, Thacker BE, Juliano MA, Pasqualini R, Arap W, Tersariol IL, Nader HB, Esko JD, Pinhal MA (2011) Inhibitory peptides of the sulfotransferase domain of the heparan sulfate enzyme, N-deacetylase-N-sulfotransferase-1. J Biol Chem 286:5338–5346.

[25] Aikawa J, Grobe K, Tsujimoto M, Esko JD (2001) Multiple isozymes of heparan sulfate/heparin GlcNAc N-deacetylase/GlcN N-sulfotransferase. J Biol Chem 276:5876–5882.

[26] Mycroft-West CJ, Abdelkarim S, Duyvesteyn HME, Gandhi NS, Skidmore MA, Owens RJ, Wu L (2024) Structural and mechanistic characterization of bifunctional heparan sulfate N-deacetylase-N-sulfotransferase 1. Nat Commun 15:1326.

[27] Kakuta Y, Sueyoshi T, Negishi M, Pedersen LC (1999) Crystal structure of the sulfotransferase domain of human heparan sulfate N-deacetylase/N-sulfotransferase 1. J Biol Chem 274:10673–10676.

[28] Carlsson P, Presto J, Spillmann D, Lindahl U, Kjellén L (2008) Heparin/heparan sulfate biosynthesis: processive formation of N-sulfated domains. J Biol Chem 283:20008–20014.

[29] Sheng J, Liu R, Xu Y, Liu J (2011) The dominating role of N-deacetylase/N-sulfotransferase 1 in forming domain structures in heparan sulfate. J Biol Chem 286:19768–19776.

[30] Tkachyova I, Fan X, LamHonWah AM, Fedyshyn B, Tein I, Mahuran DJ, Schulze A (2016) NDST1 preferred promoter confirmation and identification of corresponding transcriptional inhibitors as substrate reduction agents. PLoS One 11:e0162145.

[31] Ringvall M, Ledin J, Holmborn K, van Kuppevelt T, Ellin F, Eriksson I, Olofsson AM, Kjellén L, Forsberg E (2000) Defective heparan sulfate biosynthesis and neonatal lethality in mice lacking N-deacetylase/N-sulfotransferase-1. J Biol Chem 275:25926–25930.

[32] Fan G, Xiao L, Cheng L, Wang X, Sun B, Hu G (2000) Targeted disruption of NDST-1 gene leads to pulmonary hypoplasia and neonatal respiratory distress in mice. FEBS Lett 467:7–11.

[33] Grobe K, Inatani M, Pallerla SR, Castagnola J, Yamaguchi Y, Esko JD (2005) Cerebral hypoplasia and craniofacial defects in mice lacking heparan sulfate Ndst1 gene function. Development 132:3777–3786.

[34] Berman HM, Westbrook J, Feng Z, Gilliland G, Bhat TN, Weissig H, Shindyalov IN, Bourne PE (2000) The Protein Data Bank. Nucleic Acids Res 28:235–242.

[35] Abramson J, Adler J, Dunger J, Evans R, Green T, Pritzel A, Ronneberger O, Willmore L, Ballard AJ, Bambrick J, Bodenstein SW, Evans DA, Hung CC, O’Neill M, Reiman D, Tunyasuvunakool K, Wu Z, Žemgulytė A, Arvaniti E, Beattie C, Bertolli O, Bridgland A, Cherepanov A, Congreve M, Cowen-Rivers AI, Cowie A, Figurnov M, Fuchs FB, Gladman H, Jain R, Khan YA, Low CMR, Perlin K, Potapenko A, Savy P, Singh S, Stecula A, Thillaisundaram A, Tong C, Yakneen S, Zhong ED, Zielinski M, Žídek A, Bapst V, Kohli P, Jaderberg M, Hassabis D, Jumper JM (2024) Accurate structure prediction of biomolecular interactions with AlphaFold 3. Nature 630:493–500.

[36] Eastman P, Swails J, Chodera JD, McGibbon RT, Zhao Y, Beauchamp KA, Wang LP, Simmonett AC, Harrigan MP, Stern CD, Wiewiora RP, Brooks BR, Pande VS (2017) OpenMM 7: rapid development of high performance algorithms for molecular dynamics. PLoS Comput Biol 13:e1005659.

[37] O’Boyle NM, Banck M, James CA, Morley C, Vandermeersch T, Hutchison GR (2011) Open Babel: an open chemical toolbox. J Cheminform 3:33.

[38] Volkamer A, Kuhn D, Rippmann F, Rarey M (2012) DoGSiteScorer: a web server for automatic binding site prediction, analysis and druggability assessment. Bioinformatics 28:2074–2075.

[39] Irwin JJ, Tang KG, Young J, Dandarchuluun C, Wong BR, Khurelbaatar M, Moroz YS, Mayfield J, Sayle RA (2020) ZINC20 — a free ultralarge-scale chemical database for ligand discovery. J Chem Inf Model 60:6065–6073.

[40] Trott O, Olson AJ (2010) AutoDock Vina: improving the speed and accuracy of docking with a new scoring function, efficient optimization, and multithreading. J Comput Chem 31:455–461.

[41] Eberhardt J, Santos-Martins D, Tillack AF, Forli S (2021) AutoDock Vina 1.2.0: new docking methods, expanded force field, and Python bindings. J Chem Inf Model 61:3891–3898.

[42] Daina A, Michielin O, Zoete V (2017) SwissADME: a free web tool to evaluate pharmacokinetics, drug-likeness and medicinal chemistry friendliness of small molecules. Sci Rep 7:42717.

[43] Daina A, Zoete V (2016) A BOILED-Egg to predict gastrointestinal absorption and brain penetration of small molecules. ChemMedChem 11:1117–1121.

[44] Baell JB, Holloway GA (2010) New substructure filters for removal of pan assay interference compounds (PAINS) from screening libraries and for their exclusion in bioassays. J Med Chem 53:2719–2740.

[45] Baell JB, Nissink JWM (2018) Seven year itch: pan-assay interference compounds (PAINS) in 2017 — utility and limitations. ACS Chem Biol 13:36–44.

[46] Brenk R, Schipani A, James D, Krasowski A, Gilbert IH, Frearson J, Wyatt PG (2008) Lessons learnt from assembling screening libraries for drug discovery for neglected diseases. ChemMedChem 3:435–444.

[47] Lipinski CA, Lombardo F, Dominy BW, Feeney PJ (2001) Experimental and computational approaches to estimate solubility and permeability in drug discovery and development settings. Adv Drug Deliv Rev 46:3–26.

[48] Adasme MF, Linnemann KL, Bolz SN, Kaiser F, Salentin S, Haupt VJ, Schroeder M (2021) PLIP 2021: expanding the scope of the protein–ligand interaction profiler to DNA and RNA. Nucleic Acids Res 49:W530–W534.

[49] Abraham MJ, Murtola T, Schulz R, Páll S, Smith JC, Hess B, Lindahl E (2015) GROMACS: high performance molecular simulations through multi-level parallelism from laptops to supercomputers. SoftwareX 1–2:19–25.

[50] Huang J, MacKerell AD Jr (2013) CHARMM36 all-atom additive protein force field: validation based on comparison to NMR data. J Comput Chem 34:2135–2145.

[51] Vanommeslaeghe K, Hatcher E, Acharya C, Kundu S, Zhong S, Shim J, Darian E, Guvench O, Lopes P, Vorobyov I, MacKerell AD Jr (2010) CHARMM general force field: a force field for drug-like molecules compatible with the CHARMM all-atom additive biological force fields. J Comput Chem 31:671–690.

[52] Jorgensen WL, Chandrasekhar J, Madura JD, Impey RW, Klein ML (1983) Comparison of simple potential functions for simulating liquid water. J Chem Phys 79:926–935.

[53] Wohlwend J, Corso G, Passaro S, Getz N, Reveiz M, Leidal K, Swiderski W, Atkinson L, Portnoi T, Chinn I, Silterra J, Jaakkola T, Barzilay R (2024) Boltz-1: democratizing biomolecular interaction modeling. bioRxiv 2024.11.19.624167.

[54] Hess B, Bekker H, Berendsen HJC, Fraaije JGEM (1997) LINCS: a linear constraint solver for molecular simulations. J Comput Chem 18:1463–1472.

[55] Darden T, York D, Pedersen L (1993) Particle mesh Ewald: an Nlog(N) method for Ewald sums in large systems. J Chem Phys 98:10089–10092.

[56] Bussi G, Donadio D, Parrinello M (2007) Canonical sampling through velocity rescaling. J Chem Phys 126:014101.

[57] Parrinello M, Rahman A (1981) Polymorphic transitions in single crystals: a new molecular dynamics method. J Appl Phys 52:7182– 7190.

[58] Valdés-Tresanco MS, Valdés-Tresanco ME, Valiente PA, Moreno E (2021) gmx_MMPBSA: a new tool to perform end-state free energy calculations with GROMACS. J Chem Theory Comput 17:6281–6291.

[59] Miller BR 3rd, McGee TD Jr, Swails JM, Homeyer N, Gohlke H, Roitberg AE (2012) MMPBSA.py: an efficient program for end-state free energy calculations. J Chem Theory Comput 8:3314–3321.

[60] Watson JL, Juergens D, Bennett NR, Trippe BL, Yim J, Eisenach HE, Ahern W, Borst AJ, Ragotte RJ, Milles LF, Wicky BIM, Hanikel N, Pellock SJ, Courbet A, Sheffler W, Wang J, Venkatesh P, Sappington I, Torres SV, Lauko A, De Bortoli V, Mathieu E, Ovchinnikov S, Barzilay R, Jaakkola TS, DiMaio F, Baek M, Baker D (2023) De novo design of protein structure and function with RFdiffusion. Nature 620:1089–1100.

[61] Dauparas J, Anishchenko I, Bennett N, Bai H, Ragotte RJ, Milles LF, Wicky BIM, Courbet A, de Haas RJ, Bethel N, Leung PJY, Huddy TF, Pellock S, Tischer D, Chan F, Koepnick B, Nguyen H, Kang A, Sankaran B, Bera AK, King NP, Baker D (2022) Robust deep learning-based protein sequence design using ProteinMPNN. Science 378:49–56.

