## Supplementary Figures and Tables for "NDST1 as a substrate-reduction target in Mucopolysaccharidosis type IIIC: virtual screening, microsecond molecular dynamics, and peptide design"

**a**

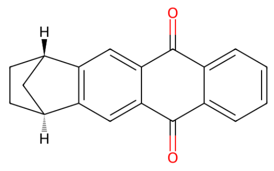

Class 1

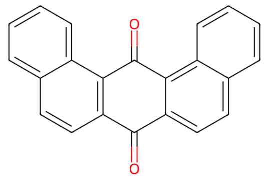

Class 2

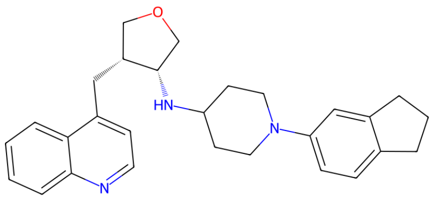

Class 3

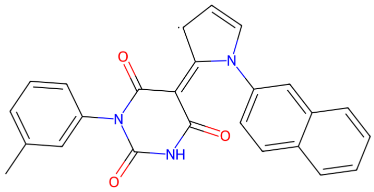

Class 4

**b**

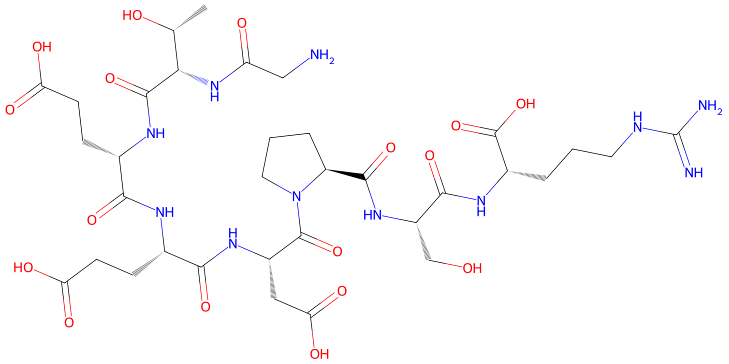

Figure S1: Chemical structures of the small-molecule and peptide binders identified in this study. **a** The four lead small-molecule scaffolds from the drug-like library (cf. Table 1 of the main text for SMILES and docking scores). **b** Linear Peptide 2 (GTEEDPSR), the retained peptide binder.

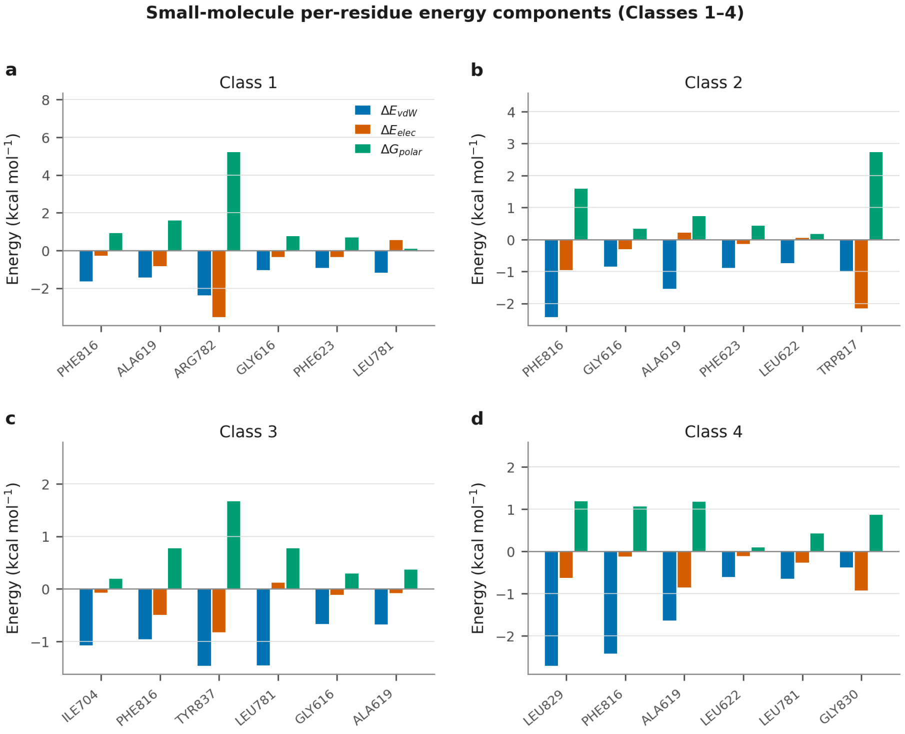


Figure S2: Van der Waals, electrostatic and polar-solvation components of the top stabilising residues for each small-molecule scaffold (**a**–**d**, Classes 1–4; cf. Fig. 7 of the main text).

Table S1: Predicted physicochemical and ADMET properties of the four lead scaffolds (SwissADME)

| Class | MW (g mol$^{-1}$) | TPSA (Å$^2$) | Rotatable bonds | HBA | HBD | log P | GI absorption | BBB |
|---|---|---|---|---|---|---|---|---|
| 1 | 274 | 34.1 | 0 | 2 | 0 | 3.16 | High | Permeant |
| 2 | 308 | 34.1 | 0 | 2 | 0 | 4.41 | High | Permeant |
| 3 | 428 | 37.4 | 5 | 3 | 1 | 4.45 | Moderate | Permeant |
| 4 | 421 | 71.4 | 3 | 3 | 1 | 3.66 | Reduced | Permeant |

Table S2: Stability metrics for the three peptide–NDST1 complexes (Linear Peptide 2 over its full 1 $\mu$s trajectory; Linear Peptide 1 and Cyclic Peptide 1 over 250 ns)

| Design | Type | Sequence | Trajectory | Peptide C$\alpha$ RMSF (Å) | Receptor C$\alpha$ RMSF (Å) | Receptor $R_g$ (Å) |
|---|---|---|---|---|---|---|
| Linear Peptide 1 | Linear | KVWEGIEKRE | 250 ns | $40.94 \pm 0.71$ | 2.56 | $32.94 \pm 0.43$ |
| Cyclic Peptide 1 | Cyclic | LVERRKED | 250 ns | $20.9 \pm 0.42$ | 1.78 | $32.67 \pm 0.27$ |
| Linear Peptide 2 | Linear | GTEEDPSR | 1 $\mu$s | $1.97 \pm 0.18$ | 2.24 | $32.92 \pm 0.39$ |
